# Neuroprotective and Mitoprotective Effects of Lemon IntegroPectin on SH-SY5Y Cells

**DOI:** 10.1101/2021.02.09.430380

**Authors:** Domenico Nuzzo, Pasquale Picone, Costanza Giardina, Miriana Scordino, Giuseppa Mudò, Mario Pagliaro, Antonino Scurria, Francesco Meneguzzo, Laura M. Ilharco, Alexandra Fidalgo, Alessandro Presentato, Rosa Alduina, Rosaria Ciriminna, Valentina Di Liberto

**Affiliations:** Istituto per la Ricerca e I’Innovazione Biomedica, CNR, via U. La Malfa 153, 90146 Palermo, Italy; Dipartimento di Scienze e Tecnologie Biologiche Chimiche e Farmaceutiche, Università di Palermo, viale delle Scienze, 90128 Palermo, Italy; Dipartimento di Biomedicina, Neuroscienze e Diagnostica Avanzata, Università di Palermo, 90134 Palermo, Italy; Istituto per lo Studio dei Materiali Nanostrutturati, CNR, via U. La Malfa 153, 90146 Palermo, Italy; Istituto per la Bioeconomia, CNR, via Madonna del Piano 10, 50019 Sesto Fiorentino, FI, Italy; Centro de Química-Física Molecular and IN-Institute of Nanoscience and Nanotechnology, Instituto Superior Técnico, Universidade de Lisboa, Av. Rovisco Pais 1, 1049-001 Lisboa, Portugal

**Keywords:** Pectin, hesperidin, IntegroPectin, neurological disease, antioxidant, neuroprotective

## Abstract

Lemon IntegroPectin obtained via hydrodynamic cavitation of organic lemon processing waste in water shows significant neuroprotective activity *in vitro*, as first reported in this study investigating the effects of both lemon IntegroPectin and commercial citrus pectin on cell viability, cell morphology, reactive oxygen species (ROS) production and mitochondria perturbation induced by treatment of neuronal SH-SY5Y human cells with H_2_O_2_. Mediated by ROS including H_2_O_2_ and its derivatives, oxidative stress alters numerous cellular processes, including mitochondrial regulation and cell signaling, propagating cellular injury that leads to incurable neurodegenerative diseases. These results, and the absence of toxicity of this new pectic substance rich in adsorbed flavonoids and terpenes, support further investigations to verify its activity in preventing, retarding, or even curing neurological diseases.

## 1. Introduction

Recently called a “universal medicine”,^[1]^ pectin is also the most valued and versatile natural hydrocolloid used by the food industry.^[2]^ Research on the broad scope biological activity of this heterogeneous, galacturonic acid-rich polysaccharide ubiquitous in plant cell walls where it acts as a cellular adhesive,^[3]^ is flourishing.^[1]^ Its antibacterial properties, for example, have been lately rediscovered.^[4]^ Lowering blood sugar levels and improving blood-sugar-related hormone function,^[5,6]^ the consumption of pectin also reduces serum cholesterol by binding with cholesterol in the digestive tract.^[7]^

Neurological disorders (ND) such as Alzheimer’s and Parkinson’s disease, and amyotrophic lateral sclerosis, causing the progressive death of neurons, are a major cause of death and disability worldwide.^[8]^ The occurrence of these morbidities has increased dramatically in recent years due to the ageing of the population in industrially developed countries, various forms of environmental pollution,^[9]^ lifestyle, nutrition (e.g., lack of sufficient intake of omega-3 essential lipids),^[10]^ and other environmental and social factors such as obesity.^[11]^ The number of patients needing neurological treatment is expected to continue to grow in the coming decades,^[8]^ along with the related healtcare costs.

Oxidative stress, the major mechanism involved in ND, alters numerous cellular processes such as mitochondrial regulation, DNA repair, and cell signaling being propagating cellular injury that leads to incurable neurodegenerative diseases.^[12]^

In 2012, Chinese scholars first reported that ginseng pectin protects neuronal cell lines from hydrogen peroxide-induced toxicity, attenuating H_2_O_2_-induced damage up to 26% in primary cortical neuron cells and human glioblastoma cells, maintaining cell integrity and decreasing nuclei condensation.^[13]^ In brief, ginseng pectin acts as a neurotrophin, protecting neurite integrity, and the team concluded that ginseng pectin might serve “as a potential therapeutic agent for neurodegenerative diseases”.^[13]^ Unfortunately, in analogy with what happened with the 1970 reports of broad scope antibacterial activity of citrus pectins,^[4]^ no subsequent report on the neuroprotective action of this and other pectins was described in the following decade.

Along with oxidative stress, inflammation of glial cells and subsequent release of neurotoxic proinflammatory molecules is an amplifier of neurological disorders,^[14]^ whereas mitochondrial dysfunction caused by amyloid β peptide is also an important component in the insurgence of Alzheimer’s disease.^[15,16,17]^

From alkaloid securinine obtained from the root of *Securinega suffruticosa*^[18]^ through citrus flavonoids,^[19]^ natural products of high antioxidant potency are increasingly investigated as neuroprotective agents for the prevention, retard or reverse of neurodegenerative diseases.^[20]^ For instance, the neuroprotective mechanism of flavonoids is thought to proceed via multiple mechanisms including suppression of lipid peroxidation, inhibition of inflammatory mediators, modulation of gene expressions and improvement of the activity of antioxidant enzymes.^[19]^ Combined with regular physical exercise,^[21]^ increasing the intake of fruits containing bioactive antioxidants is increasingly suggested to delay or inhibit neurodegeneration and correlated diseases.^[22]^

In this context, the discovery of a new citrus pectin derived via hydrodynamic cavitation of citrus processing waste (the industrial residue of citrus juice extraction) carried out directly on a semi-industrial scale was reported in late 2019.^[23]^ Called “IntegroPectin”, the properties of this pectic polymer largely differ from those of commercial citrus pectin extracted from dried citrus peel using acidic hydrolysis in hot water. Containing large amounts of flavonoids and terpenes, lemon and grapefruit IntegroPectin, for instance, are broad scope bactericidal agents showing vastly enhanced antimicrobial activity when compared to commercial citrus pectin.^[24]^ Lemon IntegroPectin is also a powerful radical scavenger, with an ORAC (Oxygen Radical Absorbance Capacity) value of 122,200 *μ*mol TE/100g (compared to 16,210 *μ*mol TE/100g of black raspberry fruit) and significant cytoprotective activity when tested with human epithelial cells.^[25]^ Now, we report that lemon (*Citrus limon*) IntegroPectin exerts significant neuroprotective activity *in vitro* against H_2_O_2_-induced damage of neuronal cells, with neuroprotection likely occurring through protection of the mitochondria.

## 2. Results and Discussion

### 2.1. Effect of pectins on cell viability upon treatment with H_2_O_2_

We carried out a dose-effect investigation on SH-SY5Y cells^[26]^ viability in response to treatment with commercial citrus pectin and lemon IntegroPectin testing the following doses via the MTT cell sensitivity assay:^[27]^ 0.0l, 0.1 and 1 mg/ml for 24 h of contact. Results in Figures 1a and 1c show evidence that both pectins did not induce any significant change in cell viability. Figure 1b and Figure 1d show that continuing the pectin treatment for the subsequent 24 h at a dose of 1 mg/ml did not induce any significant change in cell viability.

**Figure 1.**
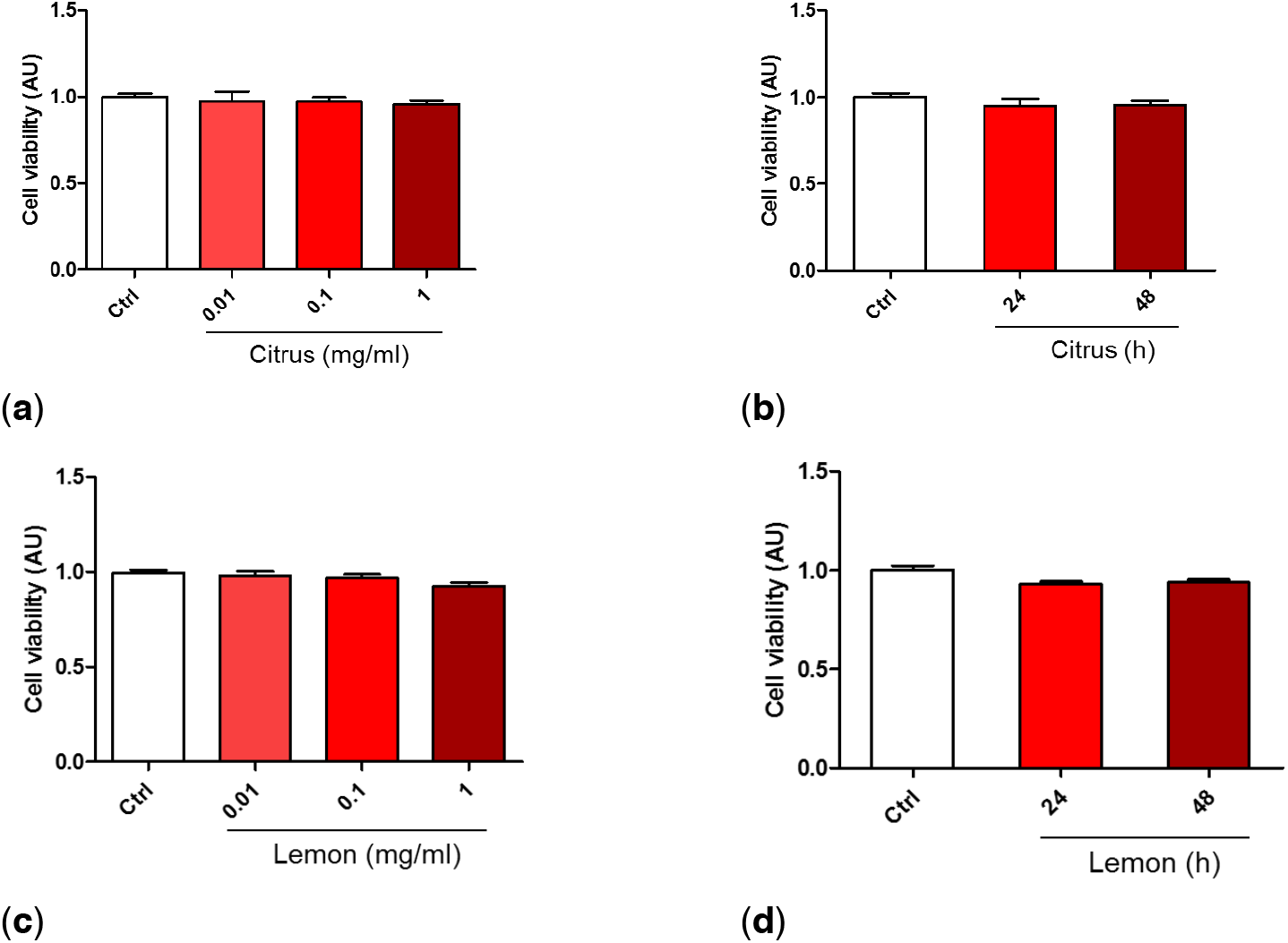
Dose and time dependent effects of commercial (citrus) and lemon IntegroPectin (lemon) pectins on neuronal cell viability: (**a**) Cell viability of citrus in dose dependent experiment; (**b**) Cell viability of citrus in time dependent experiment; (**c**) Cell viability of Lemon IntegroPectin in dose dependent experiment; (**d**) Cell viability of Lemon IntegroPectin in time dependent experiment.

We thus tested the neuroprotective effect of pectin pretreatment (Figure 2a) or co-treatment (Figure 2b). Pretreatment of citrus pectin at both 1 mg/ml and 0.1 mg/ml dose was not effective in rescuing cell viability impaired by the biocidal action of H_2_O_2_. On the contrary, lemon IntegroPectin pretreatment at the same doses induced a significant, dose-dependent increase in cell viability compared to H_2_O_2_-treated cells (Figure 2c). Similar results were obtained when pectins were co-administered with H_2_O_2_ (co-treatment,- Figure 2d). Based on these data, the pectin dose chosen for all subsequent protection experiments was 1 mg/ml.

**Figure 2.**
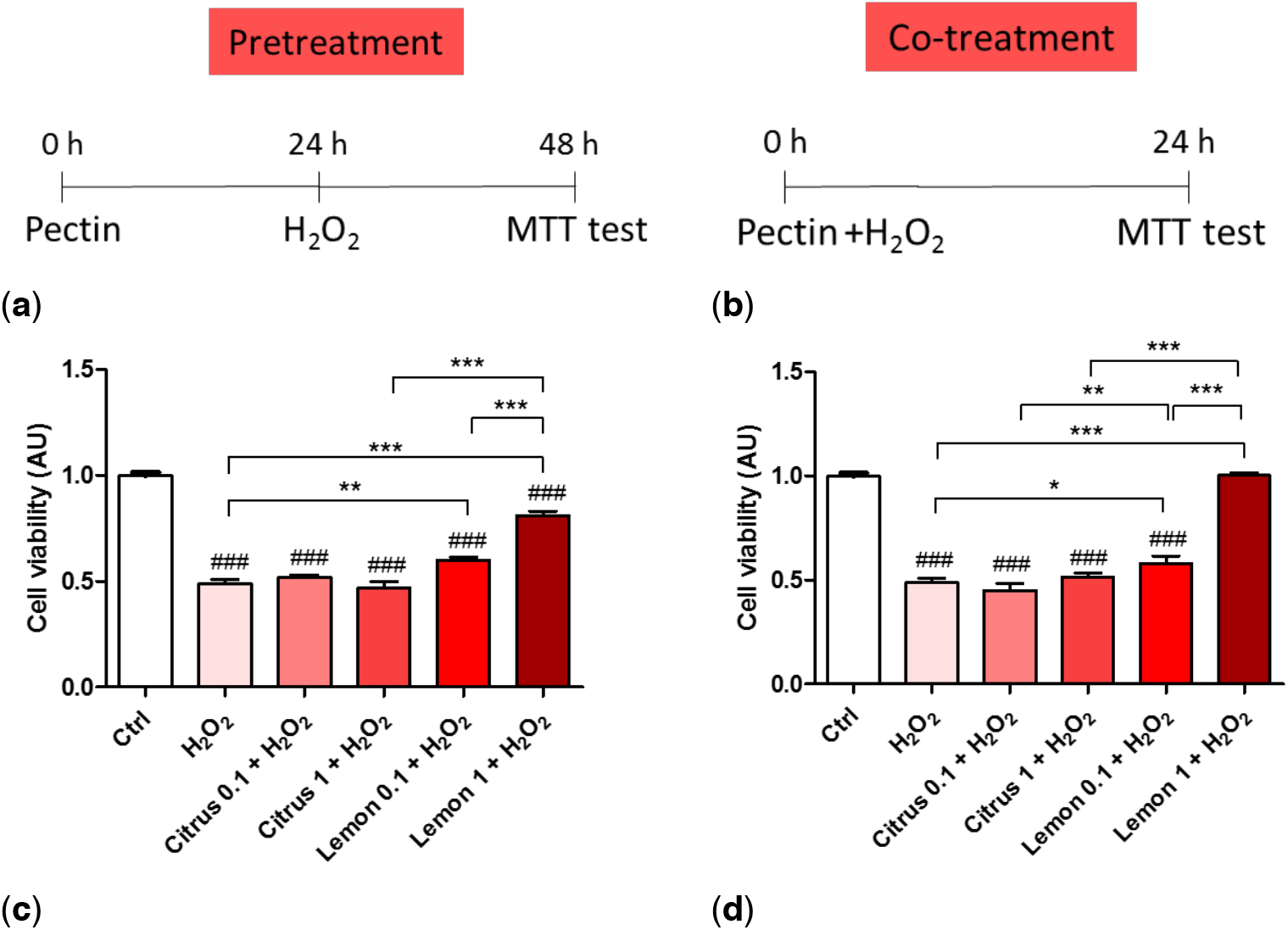
Oxidative insult during pretreatment and co-treatment whit commercial (citrus) and lemon IntegroPectin (lemon) pectins: (**a**) Scheme of cell pretreatment with pectins; (**b**) Scheme of cell co-treatment whit pectins; (**c**) Cell viability of pretreatment in dose dependent experiment; (**d**) Cell viability of co-treatment in dose dependent experiment. Tukey test: ### p < 0.001 as compared to control (Ctrl) group; * p < 0.05, ** p < 0.01, *** p < 0.001.

### 2.2 Effects of pectins on cell morphology impaired by H_2_O_2_ treatment

Since according to the MTT assay, pretreatment or co-treatment with lemon Integropectin increases cell viability, we next assessed the effect of pectin treatment on cell morphology by evaluating the cell body size (Figure 3a and 3d). Figures 3a and 3b show evidence that pretreatment of the cells with commercial citrus pectin was not able to recover cell morphology severely impacted by the H_2_O_2_ treatment. On the contrary, pretreatment of the neuronal cells with lemon IntegroPectin completely protected the neuronal cells with full retention of the cell body size (Figure 3b).

**Figure 3.**
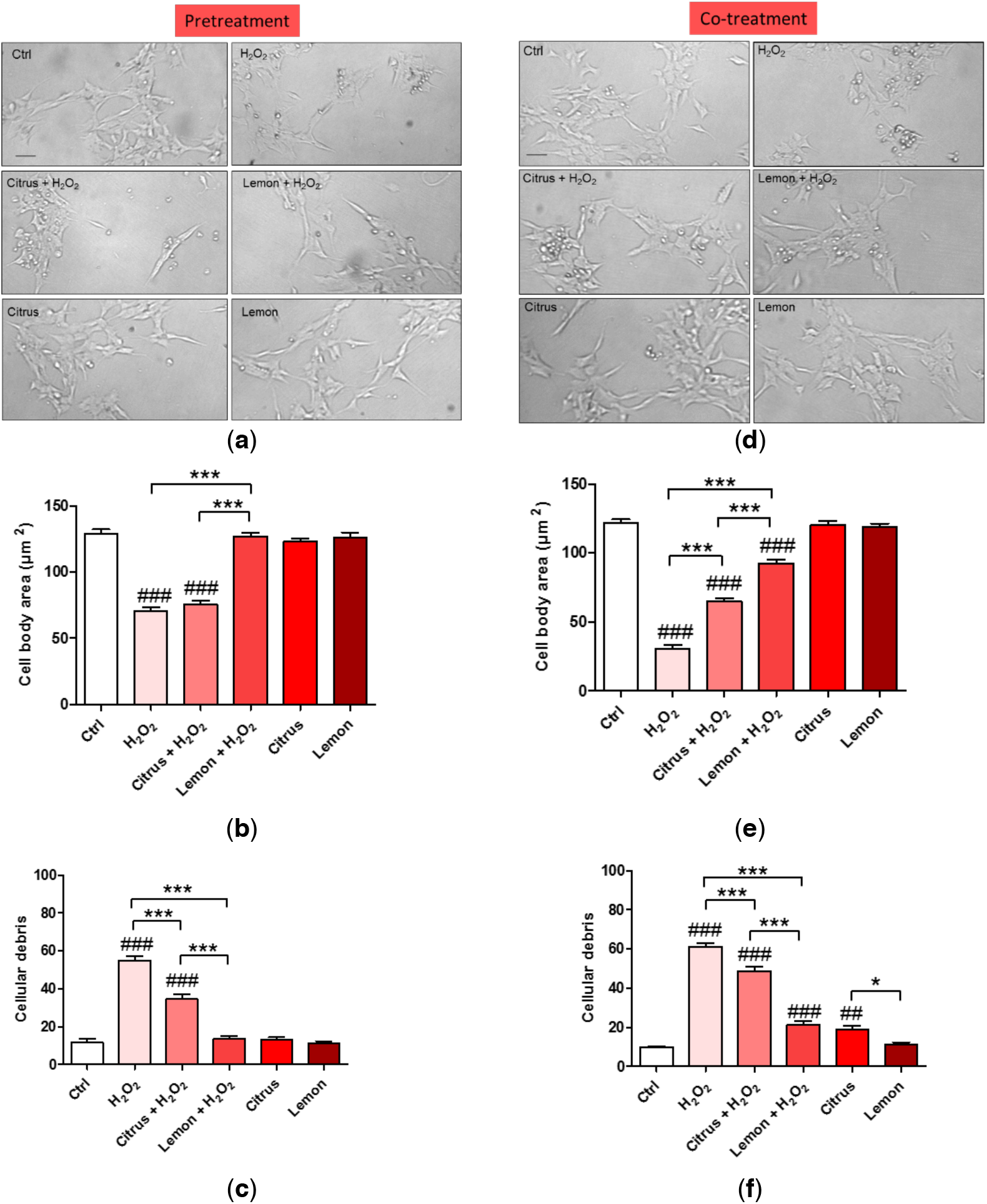
Effects of commercial (citrus) and lemon IntegroPectin (lemon) pectins on neuronal cell morphology in pretreatment and co-treatment: (**a**) Representative morphological images of the pretreatment of untreated cells (Ctrl) or treated with pectins or with H_2_O_2_ alone or in combination with pectins; (**b**) Histogram of the pretreatment of untreated cells (Ctrl) or treated or with H_2_O_2_ alone or in combination with pectins, cell body area; (**c**) Histogram of the pretreatment of untreated cells (Ctrl) or treated or with H_2_O_2_ alone or in combination with pectins, number of cells debris; (**d**) Representative morphological images of co-treatment of untreated cells (Ctrl) or treated with pectins or with H_2_O_2_ alone or in combination with pectins; (**e**) Histogram of the co-treatment of untreated cells (Ctrl) or treated or with H_2_O_2_ alone or in combination with pectins, cell body area; (**f**) Histogram of the co-treatment of untreated cells (Ctrl) or treated or with H_2_O_2_ alone or in combination with pectins, number of cells debris. Bar: 50 μm. Tukey test: ## p < 0.01, ### p < 0.001 as compared to control (Ctrl) group; * p < 0.05, *** p < 0.001.

We thus counted the number of spheroidal debris particles in each for field image, indicative of cell disintegration and death. Figure 3c shows evidence that pretreatment with lemon IntegroPectin totally prevented H_2_O_2_-induced increase in cell debris, whereas pretreatment with citrus pectin displayed only a partial neuroprotective effect.

Figures 3d and 3e show that co-treatment of the cells with commercial citrus pectin was not able to recover cell morphology severely impacted by the H_2_O_2_ treatment. On the contrary, pretreatment of the neuronal cells with lemon IntegroPectin significantly protected the neuronal cells with and rescued cell body size (Figure 3e).

Figure 3f shows that co-treatment of the neuronal cells with both pectic polymers in solution and aqueous H_2_O_2_ partially counteracted H_2_O_2_-induced increase in cell debris, with the effect of lemon IntegroPectin being significant more pronounced than that of citrus pectin. The presence of the cell debris was confirmed by the DNA cellular nuclear staining showing that treatment with H_2_O_2_ induced indeed increase in DNA damaging with degenerate nuclei being iperintense and fragmented. This condiction is reverted only by pretreatment or co-treatment with lemon IntegroPectin (Figure SIa and Figure SIb), clearly indicating a DNA protection effect of the newly discovered pectin.

### 2.3 Effects of pectins on ROS production induced by H_2_O_2_ treatment

The impact of pectins on H_2_O_2_-induced oxidative stress was assessed measuring ROS production by DCFH-DA fluorescence intensity assay. Fluorescence microscope inspection (Figure 4a) and fluorescence intensity measurement (Figure 4b) show that pretreatment of neuronal cells with lemon IntegroPectin almost completely hindered the ROS increase driven by H_2_O_2_ addition, while pretreatment with commercial citrus pectin only partially prevented it.

**Figure 4.**
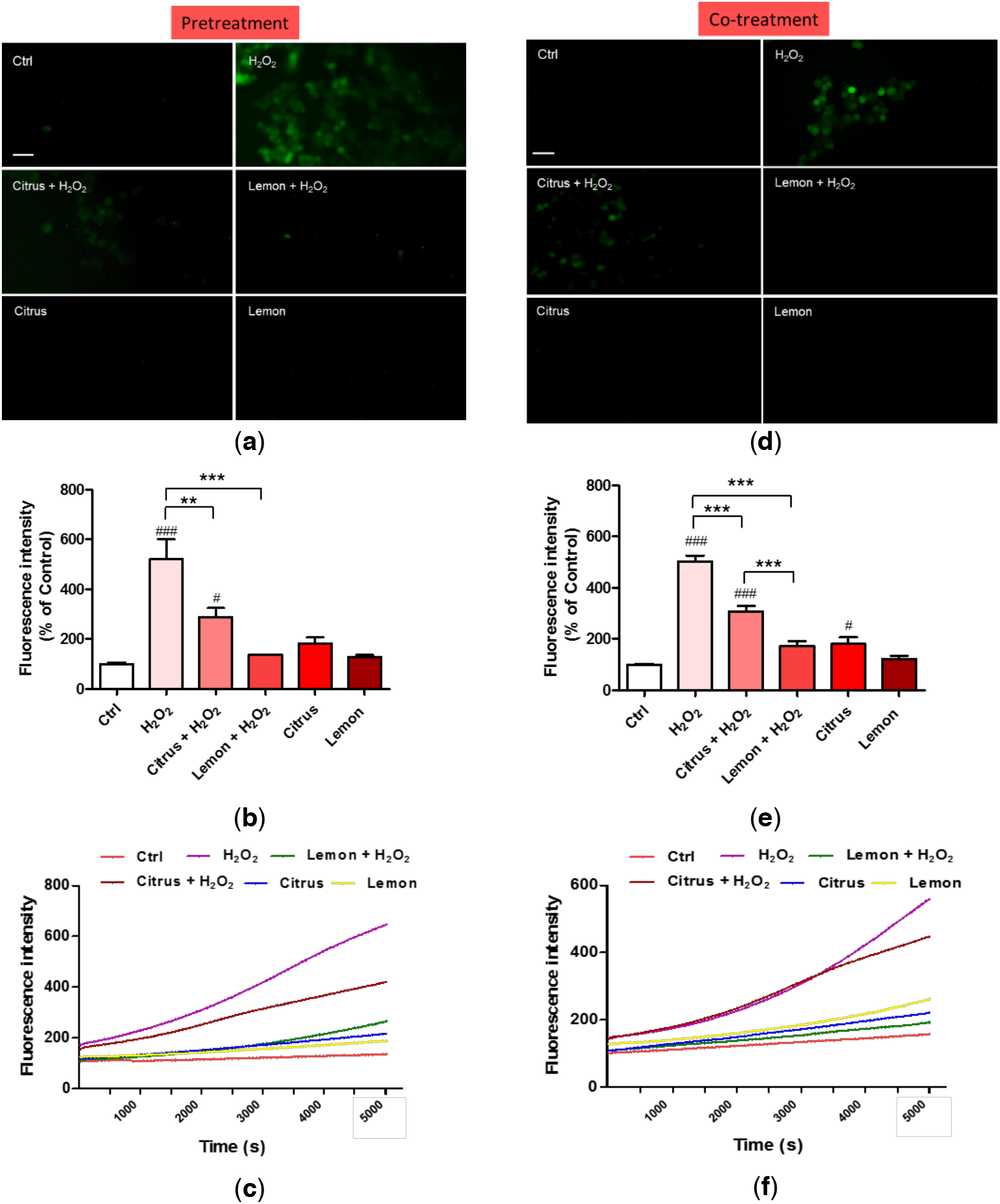
Effects of commercial (citrus) and lemon IntegroPectin (lemon) pectins on ROS production driven by exposure to aqueous H_2_O_2_: (**a**) fluorescence microscopy images of the pretreatment of untreated cells (Ctrl) or treated with pectins or with H_2_O_2_ alone or in combination with pectins; (**b**) fluorescence intensity of the pretreatment of untreated cells (Ctrl) or treated with pectins or with H_2_O_2_ alone or in combination with pectins measured by the DCFH-DA fluorescence assay; (**c**) oxidation kinetics of the pretreatment of untreated cells (Ctrl) or treated with pectins or with H_2_O_2_ alone or in combination with pectins; (**d**) fluorescence microscopy images of the co-treatment of untreated cells (Ctrl) or treated with pectins or with H_2_O_2_ alone or in combination with pectins; (**e**) fluorescence intensity of the co-treatment of untreated cells (Ctrl) or treated with pectins or with H_2_O_2_ alone or in combination with pectins measured by the DCFH-DA fluorescence assay; (**f**) oxidation kinetics of the co-treatment of untreated cells (Ctrl) or treated with pectins or with H_2_O_2_ alone or in combination with pectins. Bar: 50 μm. Tukey test: # p < 0.05, ### p < 0.001 as compared to control (Ctrl) group; ** p < 0.01, *** p < 0.001.

The kinetics of ROS production after exposure of the neuronal cells to H_2_O_2_ shows a quick and rapidly accelerating increase in ROS production (purple curve in Figure 4c). Pretreatment with commercial citrus pectin does not prevent rapid ROS accumulation, though the final amount of ROS, during the first 2 h of treatment, is lower than with H_2_O_2_ alone (brown curve in Figure 4c). Pretreatment with lemon IntegroPectin is particularly effective in lowering and delaying ROS production driven by H_2_O_2_ (green curve in Figure 4c). The ROS generation plot is linear and the growth rate is slow.

Neuronal cells co-treated with lemon IntegroPectin did not show any increase in fluorescence intensity when concomitantly exposed to aqueous H_2_O_2_ (Figures 4d and 4e). In contrast, cells co-treated with commercial citrus pectin showed a partial reduction of green fluorescence, indicative of ROS generation.

The kinetics of ROS production after exposure of the neuronal cells to H_2_O_2_ show that co-treatment of neuronal cells with lemon IntegroPectin counteracted ROS increase driven by H_2_O_2_ (green curve in Figure 4f). In contrast, co-treatment of the neuronal cells with commercial citrus pectin and hydrogen peroxide was almost completely ineffective in preventing or delaying ROS generation (purple curve in Figure 4f).

### 2.4 Effects of pectins on mitochondrial membrane potential altered by H_2_O_2_ treatment

Exacerbated ROS production damages mitochondrial components generating dysfunctional mitochondrial units. The exposure to excessive oxidative stress results in an increase in ROS concentration until a threshold level is reached that triggers the opening of the mitochondrial permeability transition pore (MPTP), leading to the collapse of the mitochondrial membrane potential and subsequent release of cytochrome C into the cytosol, which in turn initates other cellular events in the apoptotic cascade.^[28]^ Variations in the physiological mitochondrial membrane potential, an indicator of cells’ health and functional status,^[29]^ were therefore measured as changes in the accumulation of JC-1 cyanine dye red and green fluorescence signals in the cells. Mitochondrial toxin CCCP was used as a positive control. Mitochondria are sensitive to ROS, mostly generated as a by-product of cellular respiration. We briefly remind that when excited at 488 nm, JC-1 monomers emit green fluorescence with a maximum at 530 nm (green), whereas J-aggregates emit orange-red fluorescence with a maximum at 595 nm (orange-red).^[29]^ The normally green fluorescent JC-1 dye forms red fluorescent aggregates when concentrated in energized mitochondria in response to their higher membrane potential.

As displayed by Figures 5a and 5b, treatment of the neuronal cells with either commercial citrus pectin or lemon IntegroPectin did not alter the green fluorescence, indicating absence of mitotoxicity of both pectins.

**Figure 5.**
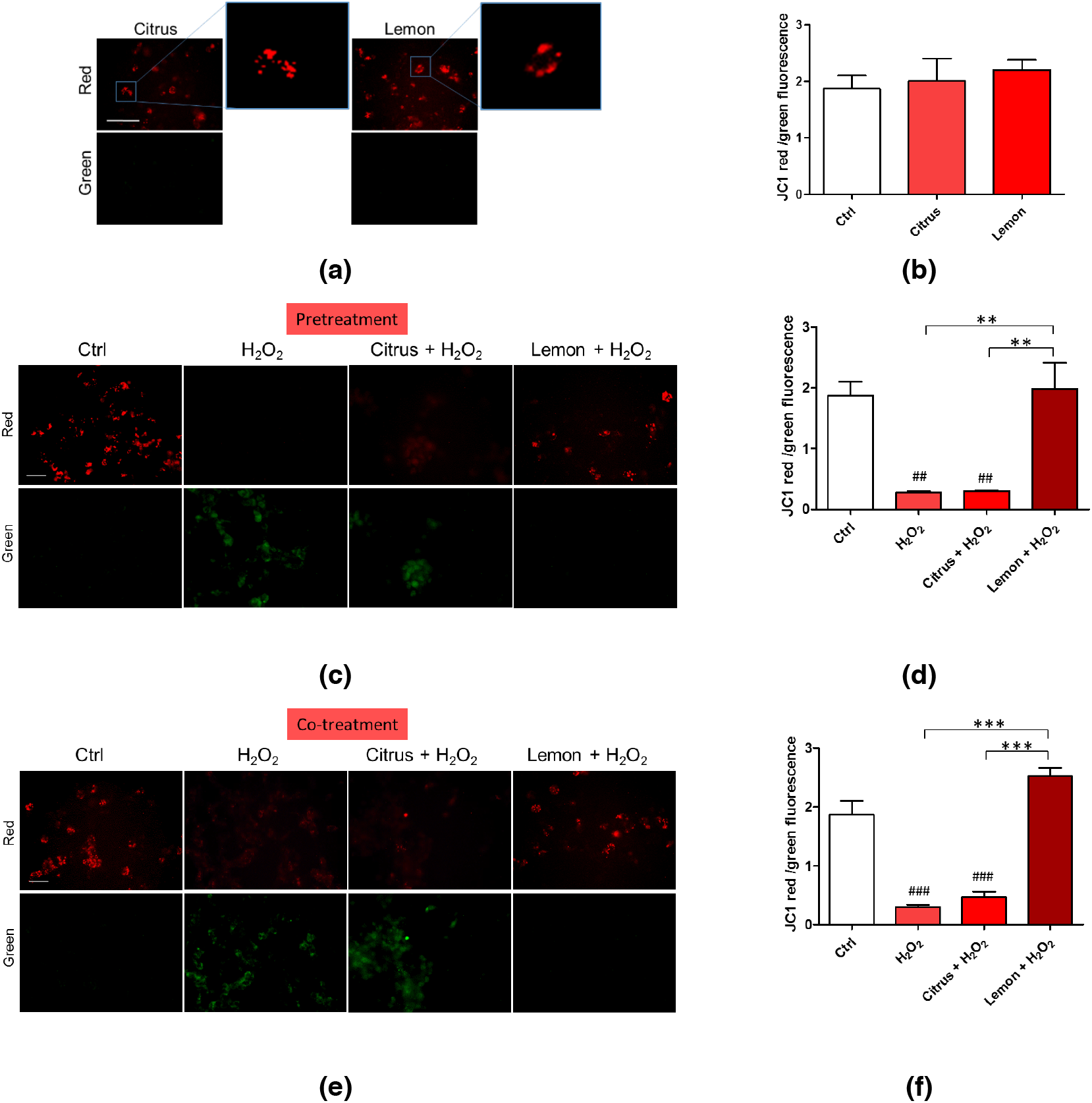
Effects of commercial (citrus) and lemon IntegroPectin (lemon) pectins on JC-1 red and green fluorescence affected by exposure to aqueous H_2_O_2_: (**a**) Fluorescence microscope inspection of cells treated with pectins alone submitted to JC-1 assay, the magnifications highlight the shape of the mitochondria, (**b**) histogram representing the ratio between red and green fluorescence intensity; (**c**) Fluorescence microscope inspection of cells untreated (Ctrl), treated with H_2_O_2_ alone, or pretreated with pectins and then with H_2_O_2_ submitted to JC-1 assay; (**d**) histogram representing the ratio between red and green fluorescence intensity in pretreatment experiment; (**e**) fluorescence microscope examination of cells untreated (Ctrl), treated with H_2_O_2_ alone or co-treated with pectins and H_2_O_2_ submitted to JC-1 assay; (**f**) histogram representing the ratio between red and green fluorescence intensity in pretreatment experiment. Bar: 100 μm.

Figures 5c and 5d show that the JC-1 red/green fluorescent signal significantly diminished following cell exposure to H_2_O_2_. Pretreatment of the cells with commercial citrus pectin and further exposed to aqueous H_2_O_2_ did not counteract the reduction of the red/green fluorescent signal. Pretreatment of the neuronal cells with lemon IntegroPectin, however, significantly reversed this effect.

Figures 5e and 5f show that co-treatment of the cells with lemon IntegroPectin and H_2_O_2_ totally counteracted the H_2_O_2_-driven reduction of the red/green fluorescent signal. On the other hand, when commercial citrus pectin was co-administered to the neuronal cells along with aqueous H_2_O_2_ it was virtually ineffective in preventing the red/green signal reduction, which turned out to be nearly identical to that driven by treatment with H_2_O_2_ alone.

### 2.5 Effects of pectins on mitochondrial morphology induced by H_2_O_2_ treatment

Changes in mitochondrial morphology occur in many neurological diseases, including Alzheimer’s,^[30]^ Parkinson’s^[31]^ and amyotrophic lateral sclerosis^[32]^ diseases. We thus analyzed mitochondrial remodeling by measuring the absorbance of isolated mitochondria. Both figures 6a and 6b show evidence that pretretment and co-treatment with lemon IntegroPectin fully counteracted the significant H_2_O_2_-induced mitochondrial remodeling. On the contrary, citrus pectin failed in producing a similar effect.

**Figure 6.**
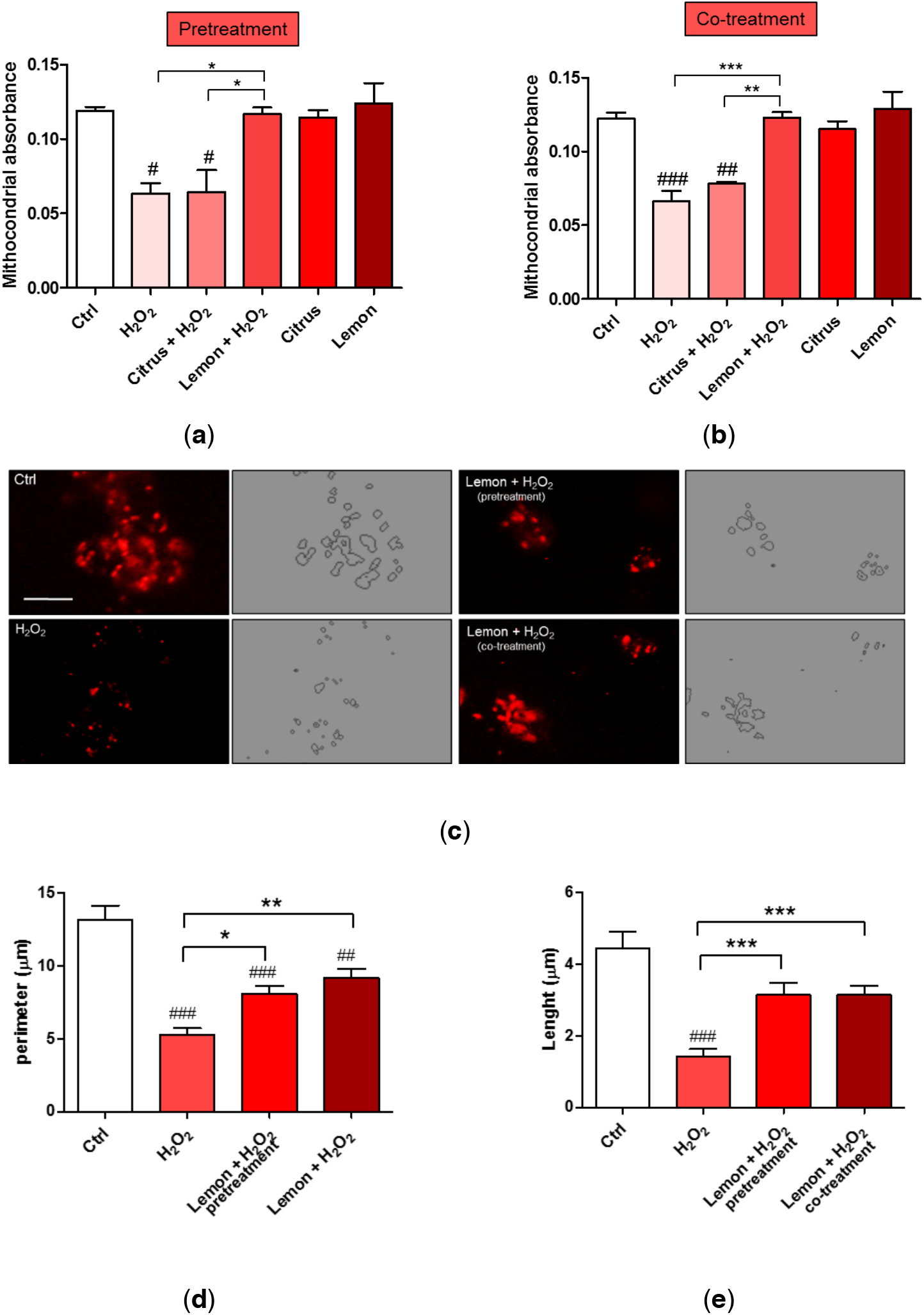
Effects of commercial (citrus) and lemon IntegroPectin (lemon) pectins on mitochondrial morphology and remodeling affected by exposure to aqueous H_2_O_2_: (**a**) Mitochondria remodeling of the pretreatment of untreated cells (Ctrl) or treated with pectins or with H_2_O_2_ alone or in combination with pectins; (**b**) Mitochondria remodeling of the co-treatment of untreated cells (Ctrl) or treated with pectins or with H_2_O_2_ alone or in combination with pectins; (**c**) Representative fluorescence microscopy images of mitochondria stained with the MitoTracker Deep Red in untreated cells (Ctrl) or treated with H_2_O_2_ alone or in combination with lemon IntegroPectin (pretreatment and co-treatment); (**d**), (**e**) Histograms representing quantification of mitochondria perimeter and length, respectively, in untreated cells (Ctrl) or treated with H_2_O_2_ alone or in combination with lemon IntegroPectin (pretreatment and co-treatment). Bar: 50 μm. Tukey test: # p < 0.05, ## p < 0.05### p < 0.001 as compared to control (Ctrl) group; * p < 0.51 ** p < 0.01, *** p < 0.001.

**Figure 7.**
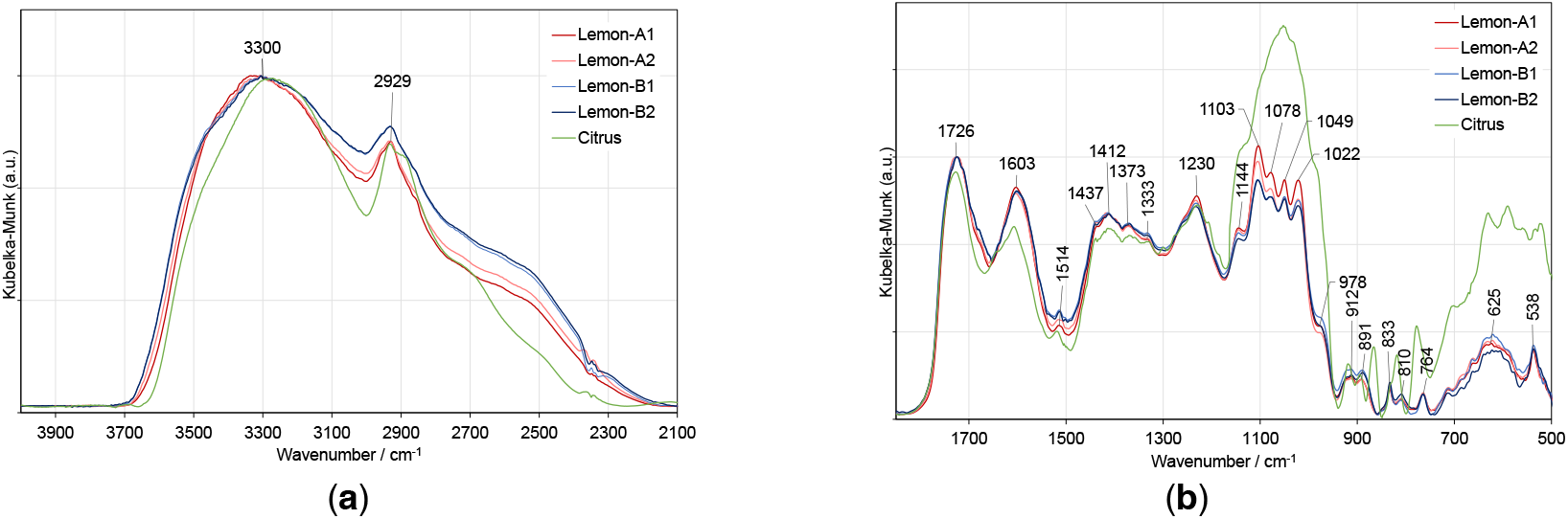
Comparison of the DRIFT spectra of lemon IntegroPectin (Lemon-A and Lemon-B) and commercial (Citrus) pectin: (**a**) 4000-2100 cm^−1^ and (**b**) 200-950 cm^−1^ spectral regions.

In light of better results obtained with lemon IntegroPectin, we analyzed its effect on the mitochondrial morphology both via cell pretreatment or co-treatment. As shown in Figure 6c, the morphology of mitochondria, stained with MitoTracker dyes in neuronal cells, during normal culture conditions is characterized by a well interconnected mitochondrial network. Treatment of the cells with aqueous H_2_O_2_ resulted in a concomitant change in mitochondrial morphology from tubular networks to fragmented puncta (circular). Remarkably, whereas a reduction in perimeter and length of the mitochondria was clearly observed after the cell treatment with hydrogen peroxide (picture in the upper left, Figure c), when lemon IntegroPectin was administered immediately before or directly along with the strong oxidizer H_2_O_2_, the morphology and mitochondrial parameters were partially recovered (Figures 6d and 6e).

### 2.6 Pectin structural insight

To interpret the outstanding properties of lemon IntegroPectin, we analyzed the molecular structure of two batches of lemon IntegroPectin (*lemon*-A and *lemon*-B). Figure 9 compares these spectra with that of a commercial (*citrus*) pectin. The 4000-2100 cm^−1^ region was normalized to thethe pyranose rings hydroxyl stretching band, and the 1850-500 cm^−1^ region was normalized to the carbonyl stretching band.

The 4000 to 2100 cm^−1^ spectral region is dominated by the broad and intense O-H stretching band of hydroxyls, overlapped with the C-H stretching band. The broad shoulder near 2550 cm^−1^, observed only in lemon IntegroPectin may be assigned to the O-H stretching vibration in free carboxyl groups.^[33]^ The two strong bands in the 1750-1500 cm^−1^ region are assigned to stretching modes of carbonyl groups from esterified *D*-GalA units and free carboxylate groups, respectively.^[34]^ The main CH_x_ and C-O-H deformation modes appear partially overlapped, in the 1500-1200 cm^−1^ region.

The five intense and partially overlapped bands observed in the 1200-950 cm^−1^ region are characteristic of the pectin backbone and side groups (Scheme 1). The band at 1144 cm^−1^ is assigned to the C-O-C stretching vibrations of the α-1,4-*D*-glycosidic bonds in the HG chains. The two strong bands at 1103 and 1022 cm^−1^ are due to skeletal stretching modes of the pyranose rings in *D*-GalA and *L*-Rha residues, present both in homogalacturonan (HG) and type I rhamnogalacturonan (RG-I) regions. The other two very intense bands, at 1078 and 1049 cm^−1^, result from neutral sugars in the side chains of RG-I and are assigned to the same stretching modes of *L*-arabinosyl and *D*-galactosyl units, respectively.^[35]^ This region is less defined and more intense in the spectrum of the commercial (*Citrus*) pectin, giving additional information on the sugar structure of these pectins. As mentioned in previous publications, it is not possible to quantify the HG content of pectin just by infrared analysis. Nevertheless, the unique shape of the cyclic ethers spectral region (1200-950 cm^−1^), in conjunction with their relatively low intensity, suggests that lemon IntegroPectin pectin has a high proportion of type I rhamnogalacturonan containing side-chains of α-1,5-*L*-arabinosyl and β-1,4-*D*-galactosyl residues when compared with the commercial (citrus) pectin.

The degree of esterification of pectin (percent of methyl-esterified carboxyl groups) was determined by the ratio of ester carboxyl to total carboxyl peak areas, estimated by decomposing the 1750-1500 cm^−1^ region into a sum of Gaussian components, using a nonlinear least-squares fitting. The main results, presented in Table 1 for the lemon Integro-Pectin replicas, yield an average degree of esterification (DM) 27±3%, much lower than that for the commercial (citrus) pectin, which is 69±2%.

**Table 1.** Computed maximum (v, cm^−1^), full width at half maximum (FWHM, cm^−1^), relative area (A, %) and assignment of the bands obtained by decomposition into a sum of Gaussian components of the 1750-1500 cm^−1^ region.

| Infrared component | $\tilde{\nu}/\text{cm}^{-1}$ | FWHM / $\text{cm}^{-1}$ | Area % |
| --- | --- | --- | --- |
| $\nu(\text{C=O})_{\text{Methyl Ester}}$ | $1743.7 \pm 0.7$ | $44 \pm 3$ | $20 \pm 2$ |
| $\nu(\text{C=O})_{\text{Methyl Ester, H.bonded}}$ | $1715.7 \pm 0.8$ | $32 \pm 2$ | $6 \pm 2$ |
| $\nu(\text{C=O})_{\text{Acid}}$ | $1692 \pm 3$ | $61 \pm 6$ | $21 \pm 4$ |
| $\nu_{\text{as}}(\text{COO}^-)$ | $1600 \pm 2$ | $101 \pm 5$ | $52 \pm 1$ |

Pectin extracted from citrus fruits is generally a high molecular weight (100 to 400 kDa) polymer, with HG proportions commonly ranging from 80 to 90%,^[36,37]^ and with a degree of methylation generally above 50%, namely high methoxy (HM) pectin. We briefly remind that pectin entitles a complex group of heteropolysaccharides containing at least eight different covalently inter-linked pectic polysaccharide types, of which HG, RG-I and, to a lesser extent, type II rhamnogalacturonan (RG-II) and/or xylogalacturonan (XGA) are the most common (Scheme 1).^[35]^

Typically, grapefruit and orange pectins extracted via conventional acid hydrolysis in hot water contain longer and more numerous RG-I polysaccharide chains than lemon pectin, affording lower intrinsic viscosity.^[37]^

The pectin industry also supplies low methoxy (LM) citrus or apple pectin (average degree of esterification DM < 50%) using enzymatic or alkaline hydrolysis under controlled conditions. HM pectin gels at pH < 3.5, in the presence of a minimum of 55% sugar, and is very unstable at pH > 5, where it will depolymerize rapidly upon heating. LM pectin tends to form gels electrostatically, stabilized by metal cations, and may gel at higher pH values. These structural characteristics and correlated properties vary according to the extraction procedure. In previous works, we have shown the advantages of hydrocavitation for improving the biological activity of citrus pectin. The essential oils (EOs) emulsify in water during the hydrocavitation of citrus processing waste (CPW), with limonene (the main citrus EO component) and linalool being converted into highly bioactive α-terpineol and terpinen-4-ol due to acid catalysis promoted by citric acid residual in the CPW enhanced by the implosion of the cavitation bubbles.^[38]^

Freeze-dried pectin obtained along with flavonoids and said emulsified citrus oil upon lyophilization contains the integral water-soluble part of waste citrus peel, including the flavonoids in a highly concentrated fashion, and constitutes a completely new biomaterial, accordingly named “IntegroPectin”.^[23,24,25,39]^ Furthermore, the DM of said citrus pectins extracted by hydrodynamic cavitation is significantly lower than 50%.

The radical scavenging action of the IntegroPectin and of commercial citrus pectin, was investigated by *in vitro* experiments using the powerful oxidant H_2_O_2_. We have also investigated the cell viability, cell morphology, ROS production induced by treatment of neuronal SH-SY5Y human cells with H_2_O_2_ in the presence of the aforementioned pectins. Moreover, for the first time, we studied the effect of pectins on the mitochondrial dysfunction, a cellular mechanism involved in different neurological diseases. The mitochondrial dysfunction driven by H_2_O_2_ was blocked by the pretreatment or co-treatment with IntegroPectin. The newly discovered IntegroPectin derived from *Citrus limon* using hydrodynamic cavitation inhibits the H_2_O_2_-induced mitochondrial membrane damage, indicating that its radical scavenging effect is also extended to mitochondrial ROS. On the other hand, little or no mitochondrial protection was exerted by commercial citrus pectin. These data indicate that only lemon IntegroPectin exerts a protective role on the cell’s central organelle, involved in several neurological dysfunctions.

We hypothesize that the neuroprotective action of lemon IntegroPectin is due to the combined action of α-terpineol, hesperidin (and likely eriotricin, too), and pectin. A neuroprotective^[40]^ and anticholinergic (cholinesterase inhibitor)^[41]^ agent, α-terpineol has been lately shown to have antidepressant-like effects.^[42]^ Hesperidin, along with eriocitrin (16.7 mg/100 ml) by far the most abundant flavonoid in lemon juice (20.5 mg/100 ml),^[43]^ exerts its neuroprotective action by improving neural growth factors and endogenous antioxidant defense functions as well as by diminishing neuro-inflammatory and apoptotic pathways.^[44]^ Hesperidin is being intensively studied also as a preventative and therapeutic agent against the development of various types of metabolic diseases.^[45]^ Lemon-derived eriotricin, known to increase antioxidant capacity and decrease inflammatory markers in mice,^[46]^ is along with hesperidin the main ingredient of a nutraceutical product recently shown to be effective, in a double-blind randomized clinical study, in managing hyperglycemia and reversal of prediabetes condition.^[47]^

Besides the neuroprotective effects of ginseng pectin,^[13]^ low-methoxyl citrus pectin was lately shown to protect pancreatic cells against diabetes-induced oxidative and inflammatory stress via galectin-3 (Gal-3), a β-galactoside-binding lectin involved in cellular inflammation and apoptosis.^[48]^ Most recently, scholars in China reported early evidence that plasma Gal-3 levels of Huntington’s disease patients correlate with disease severity, and that suppression of Gal-3 suppresses inflammation, and restores neuronal DARPP32 levels, improving motor dysfunction, increasing survival in mice affected by the disease leading the team to conclud that “Gal-3 is a novel druggable target for Huntington’s disease”.^[49]^

Pectin is a natural and specific inhibitor of Gal-3, via its multiple galactan side chains,^[50]^ and citrus pectins obtained via cavitation are particularly rich in RG-I side chains^[51]^ when compared to citrus pectins obtained via conventional acid hydrolysis of the dried citrus peel which results in the loss of most hydrophilic RG-I chains in favor of HG “smooth” regions. The RG-I regions of the pectin heteropolysaccharide, on the other hand, are preserved by using green, acid-free extraction techniques such as microwave-assisted extraction,^[52]^ acoustic,^[51]^ and hydrodynamic^[23,24,25]^ cavitation. In this regard, the low DE values observed for lemon and grapefruit IntegroPectin samples seem to indicate even greater preservation of RG-I chains through hydrodynamic cavitation in comparison with other extraction techniques, enhancing the applicative potential of these bioactive extracts.

## 3. Conclusions

In conclusion, lemon IntegroPectin, an exceptionally powerful antioxidant obtained from (organic) lemon processing waste using hydrodynamic cavitation, exerts significant *in vitro* neuroprotective and mitoprotective action on neuronal SH-SY5Y human cells treated with aqueous H_2_O_2_, a strong oxidizer involved in the cellular mechanisms leading to neurodegenerative pathologies.

The radical scavenging action of the IntegroPectin^[25]^ was confirmed by the *in vitro* experiments on neuronal cells. The H_2_O_2_-induced mitochondrial dysfunction was blocked by the pretreatment or co-treatment with IntegroPectin, indicating that its scavenging effect is also extended to mitochondrial ROS. Moreover, this newly developed pectin inhibits H_2_O_2_-induced mitochondrial membrane damage. On the other hand, little or no protection was exerted by commercial citrus pectin indicating that only lemon IntegroPectin exerts a protective role on the cell’s central organelle, involved in several pathological dysfunctions.

We ascribe these findings to the combined action of the terpenes and flavonoids adsorbed at the surface of the pectic polymer as well as to the unique structure of this new pectin of low degree of esterification and with a more pronounced abundance of RG-I “hairy” regions rich in galactose and arabinose units, capable to bind and inhibit β-galactoside-binding lectin galectin-3. As recently remarked by Sobarzo-Sánchez and co-workers suggesting the need for further clinical trials, a few clinical studies have already shown that hesperidin-enriched dietary supplements can significantly improve cerebral blood flow, cognition, and memory performance.^[44]^ In this respect, it is encouraging that lemon IntegroPectin did not show any toxic activity on the delicate neuronal SH-SY5Y cell line studied even at high doses, likewise to what happens in the treatment of human epithelial pulmonary cells.^[25]^ The effects of dietary supplementation with lemon IntegroPectin on the prevention and treatment of neurodegenerative diseases should be urgently investigated.

## 4. Experimental Section

### 4.1 Solubilization of pectins

Commercial pectin derived from citrus peels (galacturonic acid, ≥74.0%, dried basis, Sigma-Aldrich, Merck, Darmstadt, Germany), labelled “Citrus” in Figures, and lemon IntegroPectin obtained as previously described^[25]^ (labelled “Lemon” in Figures), were solubilized dissolving 1 mg of pectic polymer powder in 10 ml of Phosphate Buffer Saline (PBS, pH = 7.4, 137 mM NaCl, 2.7 mM KCl, 8 mM Na_3_PO_4_). The supernatant was collected, filtered using a 0.45 μm sartorius filter, aliquoted (1 ml/vial), and stored at +4°C.

### 4.2 Cell cultures and treatment

SH-SY5Y cells generously provided by Dr. Venera Cardile, University of Catania, Italy, were cultured in T25 tissue culture flasks. Complete Dulbecco’s Modified Eagle’s Medium and F12 (Corning, DMEM/F12; 1:1) was used, supplemented with 10% fetal bovine serum (FBS), 100 U/ml penicillin and 100 U/ml streptomycin (Sigma) and 2 mM l-glutamine in a humidified atmosphere of 95% air and 5% CO_2_ at 37 °C. The cell culture medium was every each 3 days, and the cells were sub-cultured once they reached 90% confluence.

The effects of pectins in solution on all the analyzed parameters were tested in cells cultured for 72 h in 96-wells plates. All treatments were performed at least 24 h after plating. Based on the experimental groups, the cells received the following treatments: H_2_O_2_ (200 μM for 24 h), dissolved pectins (Citrus or Lemon, 1 mg/ml, 0.1mg/ml and 0.01 mg/ml for 24 h or 48 h), a combination of pectins and H_2_O_2_, with pectins administered 24 h before (pretreatment), immediately before (co-treatment), or 3 h after H_2_O_2_ exposure (treatment). The control (Ctrl) groups received only an equal volume of the solvent medium.

### 4.3 Cell viability and cell morphology

Cells were grown at a density of 2×10^4^ cell/well on 96-well plates in a final volume of 100 μl/well. Cell viability was assessed by measuring the amount of coloured formazan by the reduction of 3-(4,5-dimethylthiazol-2-yl)-2,5-diphenyltetrazolium bromide (MTT, 0.5 mg/ml) by viable cells after 3 h incubation at 37 °C. Absorbance was measured at 570 nm with background subtraction after dissolving formazan crystals with DMSO 100 μl/well. Cell viability was expressed as arbitrary units, with the control group set to 1. For analysis of cell morphology, cells were grown at a density of 5×10^3^ cell/well on 96-well plates in a final volume of 100 μl/well. At the end of the experiments, the cells were fixed with 4% formaldehyde solution for 15 min at room temperature, washed twice with PBS, and nuclei were counterstained with the fluorescent stain 4’,6-diamidino-2-phenylindole (DAPI). The cellular images obtained using the Zeiss Axio Scope 2 microscope (Carl Zeiss, Oberkochen, Germany) were analyzed with the ZEISS-ZEN imaging software, measuring each time the cell body size and the number of cell debris *per* field.

### 4.4 Analysis of Reactive Oxygen Species

To assess ROS generation, SH-SY5Y cells were placed at a density of 1×10^4^ cells/well on 96-well plates in a final volume of 100 μl/well. At the end of the treatments, each sample was added dichlorofluorescein diacetate (DCFH-DA, 1 mM) and placed in the dark for 10 min at room temperature. After washing with PBS, the cells were analyzed by the fluorescence Zeiss Axio Scope 2 microscope (Carl Zeiss, Oberkochen, Germany) and using a Microplate Reader GloMax fluorimeter (Promega Corporation, Madison, WI 53711 USA) at the excitation wavelength of 475 nm and emission wavelength 555 nm for fluorescence intensity detection. Results were expressed as a percentage of the control group.

### 4.5 Oxidation kinetics

The oxidation kinetics was investigated by placing the treated SH-SY5Y cells at a density of 2×10^4^ cell/well on 96-well plates in a final volume of 100 μl/well. At the end of the treatment, the kinetics of ROS production was evaluated for 2 h after the addition of 2,7-dichloro-diidrofluorescineacetate (DCFH-DA, Merck, Darmstadt, Germany) using the Microplate Reader GloMax fluorimeter (Promega Corporation, Madison, WI 53711 USA) at the excitation wavelength of 475 nm and emission wavelength 555 nm.

### 4.6 Mitochondrial transmembrane potential

The mitochondrial transmembrane potential eas measured by plating and treating the cells as mentioned above. After H_2_O_2_ treatment, the cells were incubated for 30 min at 37°C with 2 mM JC-1 red dye (5,5’,6,6’-tetrachloro-1,1’,3,3’-tetraethylbenzimidazolylcarbocyanine iodide) using the MitoProbe JC-1 assay kit (Molecular Probes, USA). CCCP (carbonyl cyanide 3-chlorophenylhydrazone, 50 μM), a mitochondrial membrane potential disrupter, was used as positive control. Fluorescence emission shift of JC-1 from red (590 nm) to green (529 nm) was evaluated by the aforementioned fluorimeter and fluorescence microscope equipped with a 488 nm excitation laser.

### 4.7 Mitochondrial analysis

#### 4.7.1 Mitochondrial membrane potential analysis

For mitochondrial live imaging, cells cultured on chambers were stained with 2 μM MitoProbe JC-1 Red for 30 min at 37°C and were washed twice with growth medium. The cells were observed using the aforementioned fluorescence microscope (488 nm excitation/ 590 nm emission). Mitochondrial toxin CCCP was used as a positive control of perturbation of mitochondrial membrane potential (data not shown).^[53]^

#### 4.7.2 Mitochondrial morphology analysis

Cellular Mitochondria were stained using the MitoTracker Deep Red (Invitrogen, Carlsbad, CA, USA) dye (20 nM) for 15 min, washed twice with growth medium, and observed under the fluorescence microscope (488 nm excitation/ 590 nm emission). Original fluorescence images were converted to binary images. Mitochondrial shapes were obtained by visualizing the mitochondria outlines automatically drawn by the ImageJ software (Public Domain, BSD-2 license, National Institutes of Health, Bethesda, MD, USA). Morphometric mitochondria data, length and perimeter, were calculated from each mitochondrial outline (N 500).

#### 4.7.3 Remodeling neuron isolated mitochondria

The remodeling of neuron isolated mitochondria was evaluated according to a published protocol,^[54]^ by measuring the changes in the absorbance of the mitochondrial suspensions at 540 nm using a GloMax Discover multimode plate reader (Promega, Italy). Briefly, a volume corresponding to 20 μg of mitochondrial proteins was incubated with 50 μL of PBS pH 7.4. The absorbance was monitored for 5 min at 37°C at 540 nm, and the mitochondrial remodeling was indicated by a decrease in the absorbance.

### 4.8 Pectin structural characterization

The molecular structure of four lemon IntegroPectin samples (two replicas of two batches) and a commercial pectin (Acros Organics) was characterized by Diffuse Reflectance Infrared Fourier Transform (DRIFT) spectroscopy, using a Mattson RS1 FTIR spectrometer (Mattson Instruments, Madison, WI, USA) equipped with a wide band MCT detector and a Specac selector, in the 4000 to 500 cm^−1^ range, at 4 cm^−1^ resolution. The spectra were the result of ratioing 500 co-added single beam scans for each sample (ground pectin powder diluted in grinded FTIR grade KBr, in the appropriate proportion to assure the validity of the Kubelka-Munk assumptions) against the same number of scans for the background (grinded KBr). The spectra were converted to Kubelka-Munk units and further processed using Microsoft Excel.

### 4.9 Statistical analysis

Data analysis was performed using GraphPad Prism 8 software (GraphPad Software, La Jolla, CA, USA). The results are presented as mean ± SE, and in some cases are expressed as arbitrary units, with controls equal to 1, or as a percentage of the control value. Statistical evaluations were performed by one-way ANOVA, followed by Tukey Post-Hoc test. Differences in P-value less than 0.05 were considered statistically significant.

## Supporting Information

Appendix A (available online).

## Acknowledgments

This study is dedicated to the memory of Prof. Natale Belluardo, neuroscientist and gentleman, for all he has done at Palermo’s University. We thank OPAC Campisi (Siracusa, Sicily, Italy) for kindly providing the lemon processing waste from which the lemon IntegroPectin was obtained.

---

**Scheme 1.**
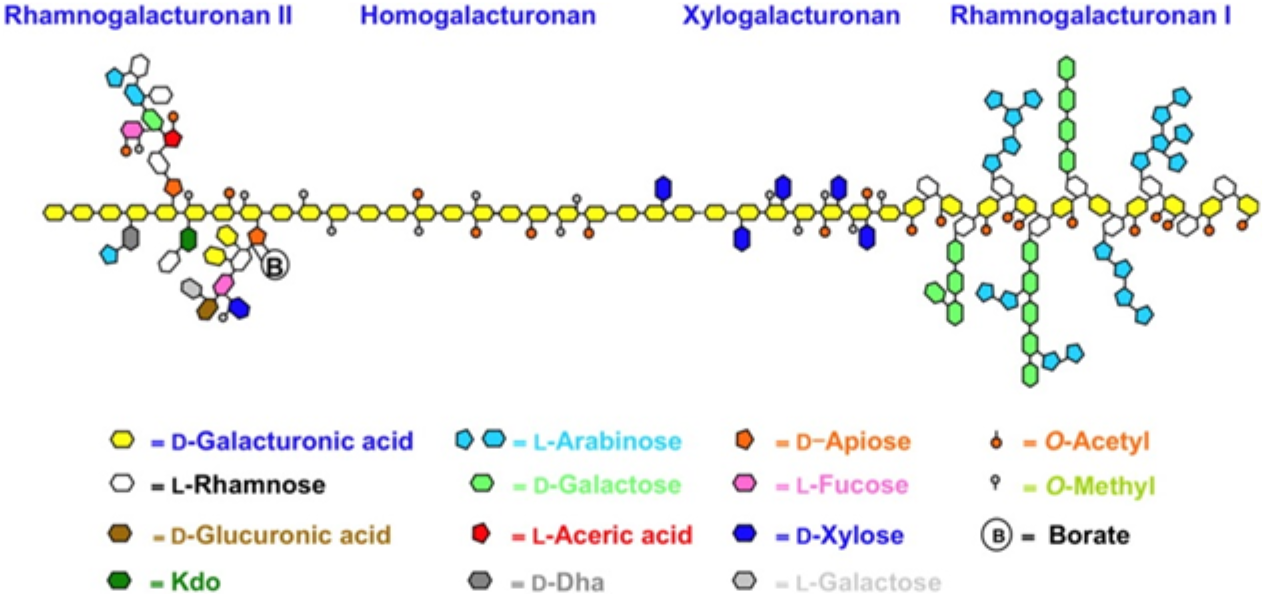
Simplified structure of pectin.

## Supplementary Information - Appendix A

**Figure S1.**
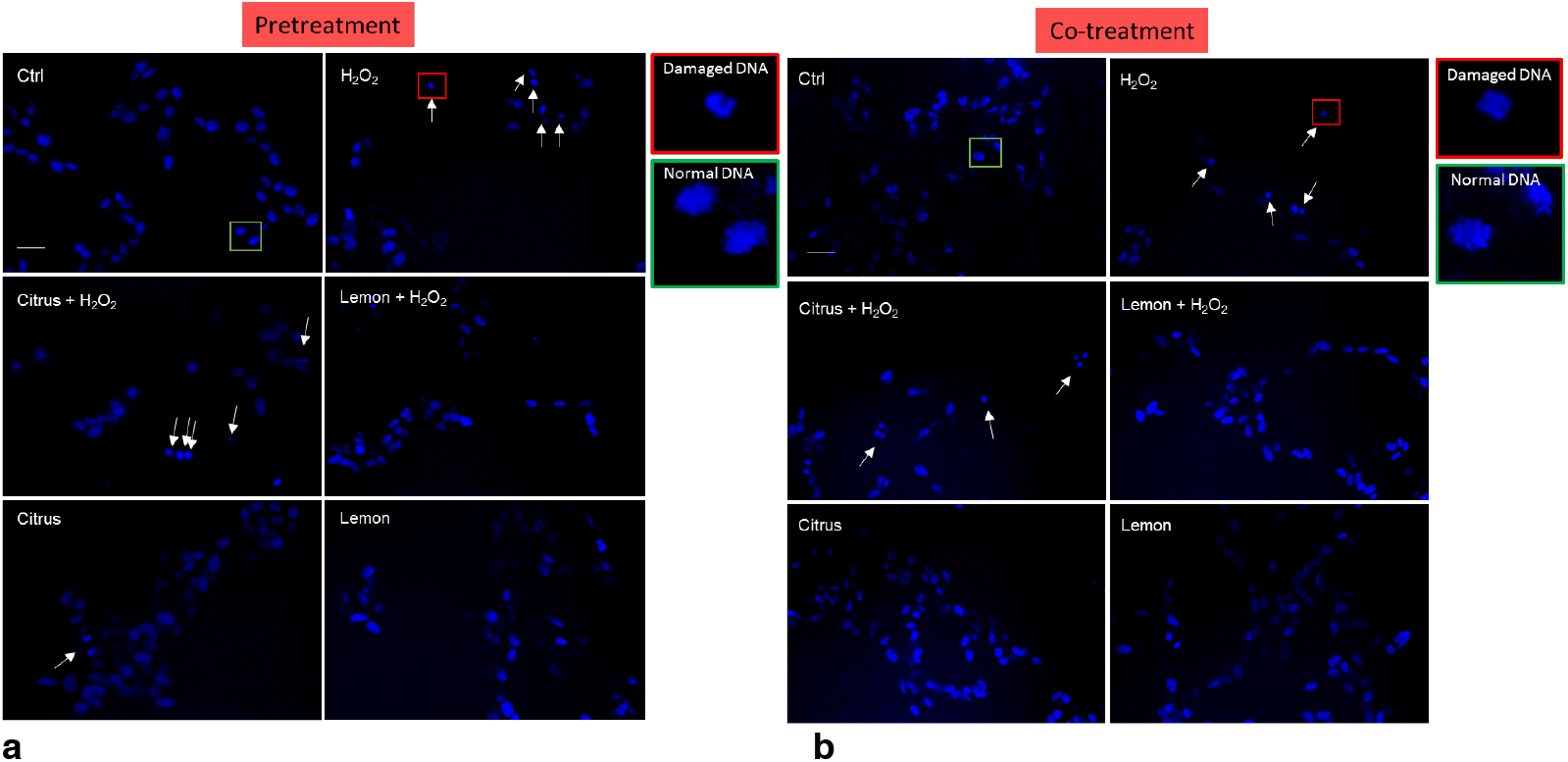
Effects of commercial (citrus) and lemon IntegroPectin (lemon) pectins on DNA damage: microscopy inspection for (**a**) pretreatment, and (**b**) co-treatment images of untreated cells (Ctrl) or treated with pectins or with H_2_O_2_ alone, or in combination with pectins. Bar: 100 μm. The square of the enlargements highlights the normal end of the damaged DNA.

